# Smoothened inhibition of PKA at cilia transduces Hedgehog signals

**DOI:** 10.1101/2025.04.01.646243

**Authors:** Thi D. Nguyen, Mia J. Konjikusic, Lorenzo M. Del Castillo, Jeremy F. Reiter

**Author notes:** Correspondence should be addressed to J.F.R.

## Abstract

Hedgehog (HH) signaling in vertebrates is dependent on the primary cilium, an organelle that scaffolds signal transduction. HH signals induce Smoothened (SMO) enrichment in the cilium and indirectly triggers the conversion of GLI proteins into transcriptional activators of HH target genes. Recently, SMO has been shown to inhibit protein kinase A (PKA). To test the hypothesis that SMO specifically inhibits PKA at cilia to activate the HH signal transduction pathway, we developed a ciliary PKA biosensor. Activation of the HH signal transduction pathway by either Sonic hedgehog (SHH) or SMO agonist (SAG) inhibited ciliary PKA activity. Blocking SMO phosphorylation by GRK2/3 prevented ciliary SMO from inhibiting ciliary PKA activity. Gα_i_ was dispensable for SMO inhibition of ciliary PKA. In contrast, mutating the SMO C-terminal tail protein kinase inhibitor (PKI) pseudosubstrate site interfered with the ability of SMO to inhibit ciliary PKA. Therefore, HH signaling is transduced via SMO direct inhibition of PKA at cilia, in a manner dependent on GRK2/3.

## Introduction

The Hedgehog (HH) signaling pathway is a critical means of cell-cell communication used by metazoans to coordinate development and homeostasis of many tissues (***1, 2***). Indispensable to vertebrate HH signaling is the primary cilium, a microtubule-based organelle that projects itself from the body of the cell (***3–5***). Though the ciliary membrane is contiguous with the plasma membrane and the cilioplasm is not membrane bound, the composition of the primary cilium is distinct from that of the rest of the cell (***6–11***). For HH signal transduction, the primary cilium acts as a specialized micro-environment in which signaling functions distinctly from elsewhere in the cell (***12–14***).

In the absence of HH signals, the HH receptor, PTCH1, localizes to the ciliary membrane and keeps the downstream signal transduction pathway off (***15***). In the presence of HH signals, HH binds to PTCH1, allowing the seven-pass transmembrane protein Smoothened (SMO) to accumulate in the ciliary membrane (***16***). Ciliary SMO is required for activation of GLI transcription factors, the effectors of HH signaling in many tissues (***17, 18***). How SMO activates GLI transcription factors remains an area of active investigation.

One possible mechanism by which SMO regulates GLI transcription factors is via protein kinase A (PKA). PKA represses HH signal transduction by phosphorylating GLI proteins to trigger the formation of their repressor forms, referred to as GLI-R (***18–20***). PKA is activated by cAMP, a second messenger mediating some forms of GPCR signaling (***21***).

Many G protein-coupled receptors (GPCRs) signal by acting as guanine nucleotide exchange factors for small GTPases including Gα_s_, Gα_i_ and Gα_o_ (***22***). These GTP-bound Gα proteins regulate the activity of adenylyl cyclases, enzymes that generate the second messenger cyclic adenosine monophosphate (cAMP) (***23***). Adenylyl cyclases are stimulated by Gα_s_ and inhibited by Gα_i/o_ (***24***). cAMP activates its principal effector, PKA (***25***). Gα_i/o_ has been investigated as an effector of SMO in HH signal transduction (***26–32***).

Beyond cAMP, some proteins, known as protein kinase inhibitors (PKIs), regulate PKA activity. PKIs directly bind to and inhibit the catalytic subunit of PKA (PKA-C) (***33***). Recent work has demonstrated that the C-terminus of SMO can bind PKA and function as a PKI (***32, 34***). We investigated whether SMO inhibits PKA at cilia to transduce HH signals, and if it does, whether PKA regulation is mediated through Gα_i/o_ or through direct inhibition of the catalytic subunit.

Another regulator of HH signal transduction is GPR161, a Gα_s_-coupled GPCR that localizes to the primary cilium in the absence of HH signals (***35–38***). A model of HH signal transduction is that GPR161, signaling through Gα_s,_ activates adenylyl cyclases to increase levels of cAMP, activating PKA, and thus triggers the formation of GLI-R (***35***).

Previously, we found that a pool of PKA localizes to the cilium and that inhibition of ciliary PKA, but not non-ciliary PKA, is sufficient to activate HH-dependent transcription (***14***). Inspired by recent discoveries that SMO inhibits PKA (***32, 34***), we hypothesized that SMO activates HH signaling by specifically inhibiting PKA at cilia. To begin to test this hypothesis, we developed a sensitive measure of ciliary PKA activity.

## Results

### Development of a biosensor of ciliary PKA activity

A previously developed biosensor of cytosolic PKA activity (**39, 40**) is based on vasodilator-stimulated phosphoprotein (VASP) Ser^157^, which is phosphorylated specifically by PKA (***41–44***). To localize this PKA-phosphorylated peptide at cilia, we fused VASP amino acids 148 to 164 to ARL13B, a ciliary protein, and GFP (**Fig. 1A**). Stable expression of ARL13B-GFP-VASP^148-163^ in a clonal NIH/3T3 cell line revealed that it, as predicted, localized to cilia (**Fig. 1B-D**).

**Figure 1:**
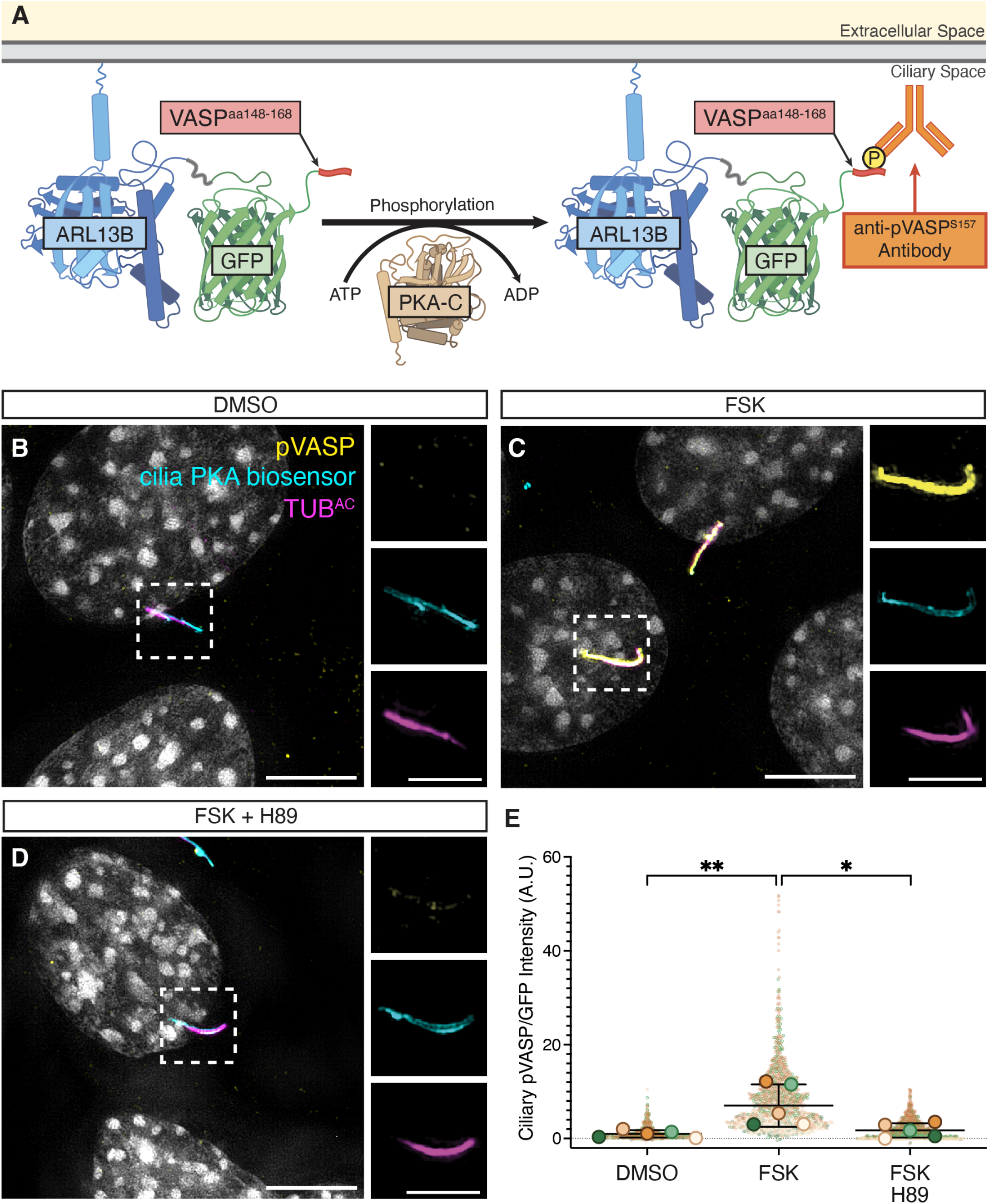
Cilia PKA biosensor detects ciliary PKA activity. (A) Schematic of ARL13B-GFP-VASP^148-163^, the cilia PKA biosensor. (**B-D**) Immunofluorescence imaging of NIH/3T3 cells stably expressing the cilia PKA biosensor. Cells were serum-starved and then treated with either DMSO (B), FSK (100nM for 15 minutes) (C), or both FSK and H89 (100nM and 20μM, respectively, for 15 minutes) (D). Cells immunostained for pVASP (pVASPS^157^, yellow), cilia PKA biosensor (GFP, cyan), cilia (acetylated tubulin, TUB^AC^, magenta) and nuclei (Hoescht, grey). Scale bars for larger images are 5μM, and for insets are 2.5μM. (**E**) Quantification of ciliary pVASP intensity normalized to ciliary GFP intensity. Representative images used for quantification are in Figure S1A-C. Each biological replicate is color coded. Significance was determined via one-way ANOVA of the means of each biological replicate, followed by Šídák’s multiple comparison test. (P values are indicated as follows: *p < 0.04, ** p < 0.003. Data are represented are means of replicates ± SD.)

To detect PKA-mediated phosphorylation of the VASP peptide, we employed a monoclonal antibody that specifically recognizes VASP phosphorylated at Ser^157^ (pVASP) (***39***). To test this detection mechanism, we treated ciliary VASP-expressing cells with the adenylyl cyclase agonist forskolin (FSK), which induces cAMP production (***45***). Immunofluorescence imaging of ciliary VASP-expressing cells using the pVASP-specific antibody revealed that the basal level of ciliary pVASP was low and dramatically increased by FSK (**Fig. 1C, 1E, S1A-B, S1D-E**). Inhibiting PKA with H89 blocked the FSK-induced increase in ciliary pVASP (**Fig. 1D-E, S1C-D**). As this assay measures ciliary PKA-dependent phosphorylation activity, we refer to ARL13B-GFP-VASP^148-163^hereafter as the cilia PKA biosensor.

We tested the dynamic range of the cilia PKA biosensor in response to different concentrations of FSK as well as different durations of FSK treatment (**Fig. S1A-E**). FSK exerted dose- and time-dependent activation of the cilia PKA biosensor. Treatment of cells with 100nM of FSK, for 15 minutes, produced a half-maximal increase in cilia PKA biosensor activity.

### Ciliary PKA activity is graded during zebrafish development

During zebrafish development, somites are patterned by SHH produced by the notochord, required for differentiation of muscle pioneers and slow muscle fibers (***46***). Because somites are generated in an anterior to posterior manner, at any developmental timepoint, the anterior somites are more mature whereas posterior somites are still differentiating and are actively responding to SHH (***47, 48***).

Therefore, we hypothesized that, if ciliary PKA activity is suppressed by HH signaling, zebrafish somites would exhibit a posterior to anterior gradient of ciliary PKA activity. To test this hypothesis, we expressed the cilia PKA biosensor in zebrafish embryos and stained for pVASP and GFP (**Fig. 2A-D**). Cilia PKA biosensor activity was higher in anterior somites and lower in posterior somites (**Fig. 2E**). Thus, in vivo, ciliary PKA active is dynamic, and active HH signaling is associated with decreased ciliary PKA.

**Figure 2:**
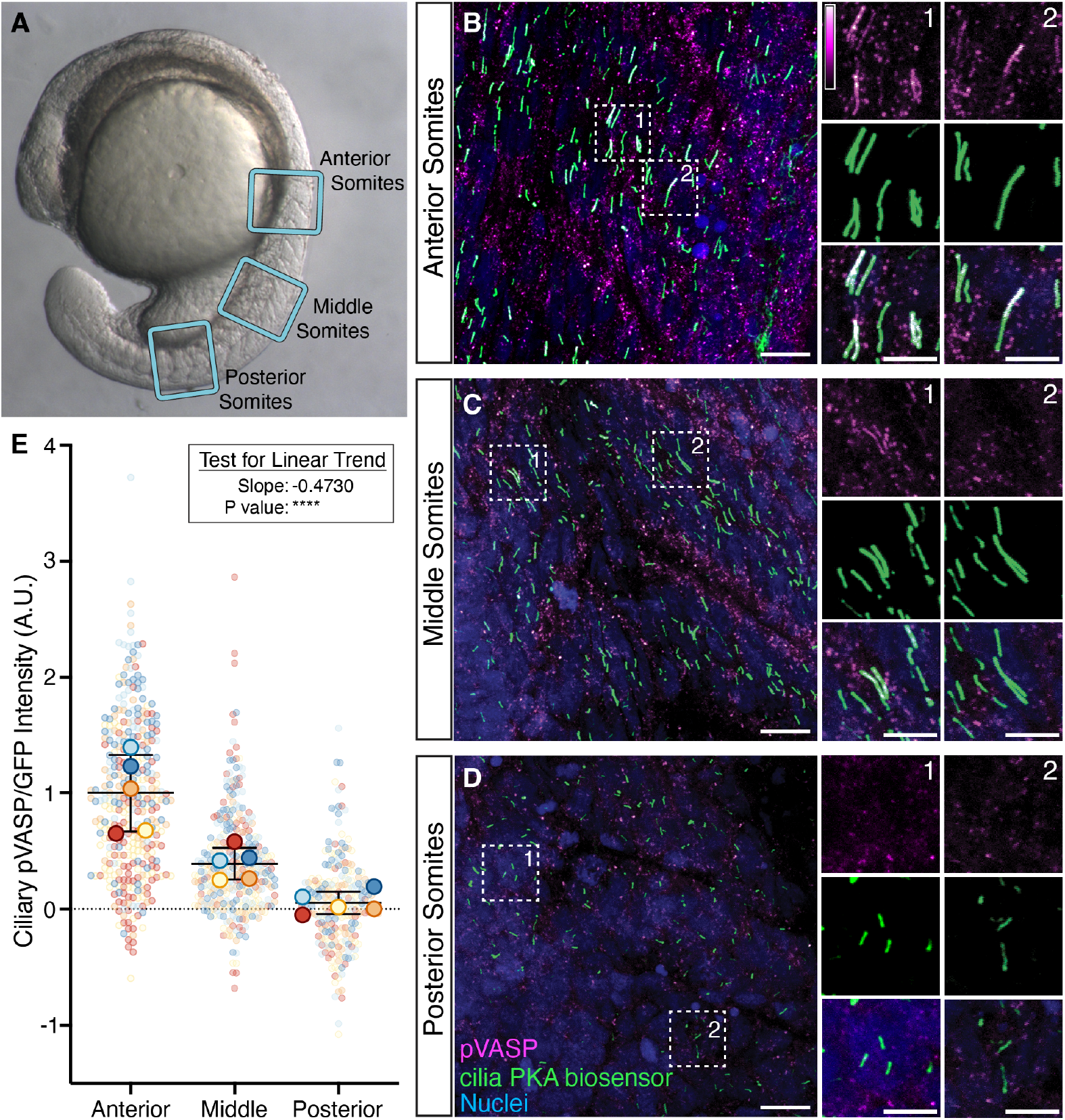
Ciliary PKA activity is graded from anterior to posterior somites in zebrafish development. (**A**) 18 somite-stage zebrafish embryo, somites 2-4 of which are designated anterior, somites 7-9 of which are designated middle, and somites 12-14 of which are designated posterior. (**B**) Immunofluorescence images of somites from zebrafish injected with 500pg mRNA encoding cilia PKA biosensor and stained for pVASP (pVASP^S157^, magenta), cilia PKA biosensor (GFP, green), and nuclei (Hoescht, blue). Scale bar, 10μM. (**C**) Quantification of ciliary pVASP intensity normalized to ciliary GFP within each somite region. Each biological replicate, one fish, is color coded. Significance was determined via one-way ANOVA of the means of each biological replicate, followed by a test for linear trend. (****p < 0.0001. Data are represented are means of replicates ± SD.)

### Active SMO suppresses ciliary PKA activity

As SMO can inhibit PKA (***34***), we hypothesized that SMO inhibits PKA at the cilium to activate HH signal transduction (**Fig. 3A**). Assays for PKA inhibition, such as with Gα_i/o_-coupled GPCRs, are typically conducted under conditions of cAMP stimulation, often with forskolin (FSK), an agonist of adenylyl cyclase (***45, 49***). Therefore, to test whether SMO inhibits ciliary PKA, we treated cilia PKA biosensor-expressing NIH/3T3 cells with FSK, FSK and H89, or FSK and Smoothened Agonist (SAG) (***50, 51***) (**Fig. 3B-G**). SAG, like H89, blocked FSK-mediated activation of the cilia PKA biosensor (**Fig. 3C-F**). Treating cilia PKA biosensor-expressing cells with FSK and SHH revealed that, like SAG, SHH blocked FSK-mediated activation of the cilia PKA biosensor (**Fig. 3F**). Thus, activating HH signal transduction blocks PKA activity in the primary cilium.

**Figure 3:**
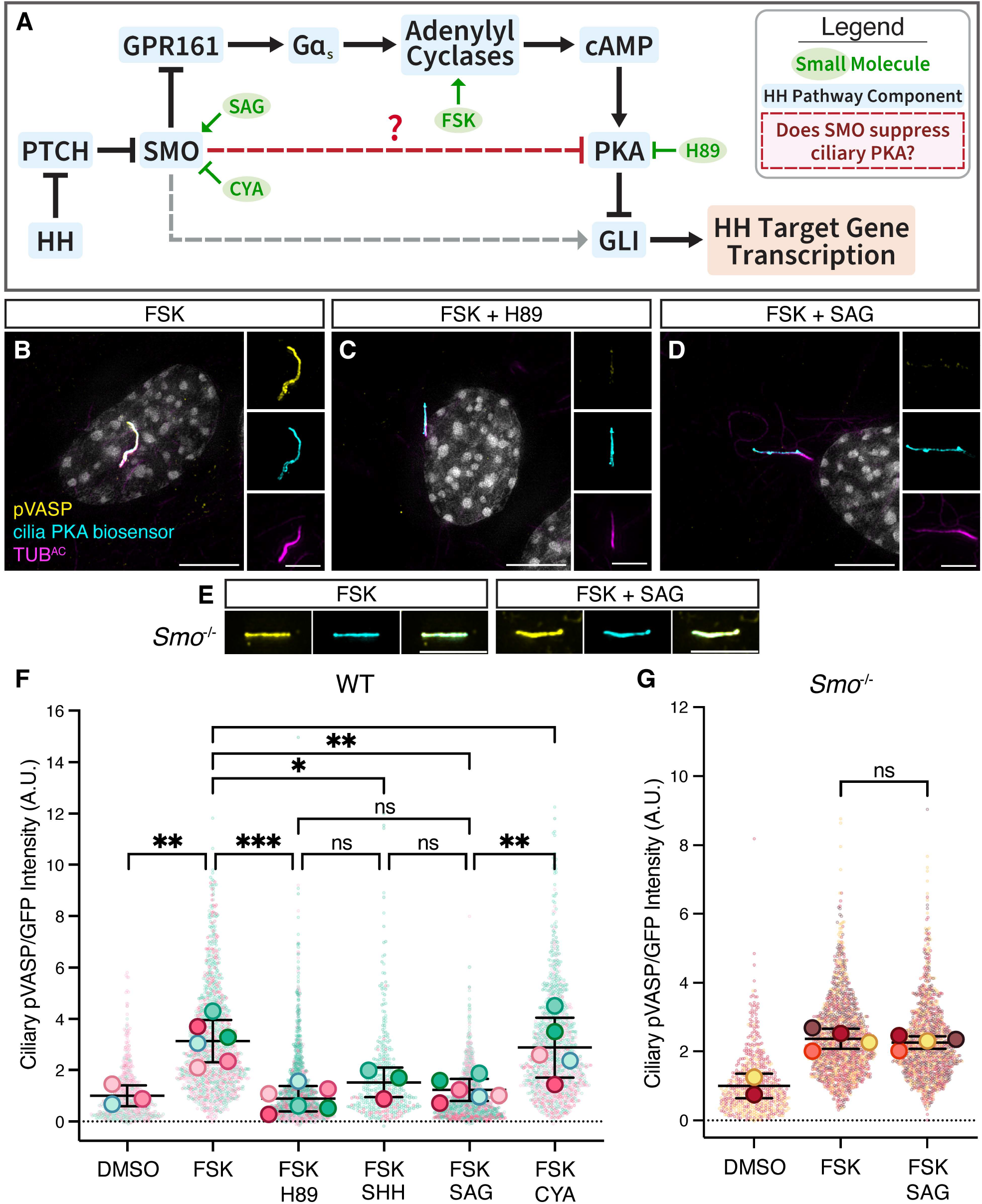
Active SMO inhibits PKA activity at cilia. (**A**) Schematic of a working model of HH signal transduction. Red line indicates part of the system investigated in this figure. (**B-E**) Immunofluorescence images of cilia PKA biosensor cells or *Smo*^-/-^ cilia PKA biosensor cells serum-starved and treated with either FSK (100nM for 15 minutes) (B), FSK and H89 (100nM and 20μM, respectively, for 15 minutes) (C), or SAG and FSK (100nM for 24 h and 100nM for 15 minutes, respectively) (D,E). Images depict cells stained for pVASP (pVASP^S157^, yellow), cilia PKA biosensor (GFP, cyan), cilia (TUB^AC^, magenta), and nuclei (Hoescht, grey). Scale bars for larger images, 5 μM (B-E). Scale bars for insets are 2.5 μM (B-D). (**F**) Quantification of ciliary pVASP intensity in cilia PKA biosensor cells. Cells in the FSK and SHH condition were treated with 24 hours of 4nM SHH followed by 100nM FSK for 15 minutes. Cells in the FSK and CYA condition were treated with 24 hours of 5μM cyclopamine followed by 100nM FSK for 15 minutes. Some datapoints for this panel are part of a dataset also used in Figure 4. (**G**) As with F, but with quantification of ciliary pVASP intensity in *Smo*^-/-^ cilia PKA biosensor cells. Data for this panel is also used in Fig. 6C and S6.For all plots, each biological replicate is color coded. Significance was determined via one-way ANOVA of the means of each biological replicate, followed by Šídák’s multiple comparison test. P values are indicated as follows: *p < 0.04, ** p < 0.003 and ***p < 0.0002. Data are represented are means of replicates ± SD.

To test if HH pathway-mediated inhibition of ciliary PKA is mediated by SMO, we used clustered regularly interspaced short palindromic repeats (CRISPR)–mediated editing to inactivate Smo in NIH/3T3 cells. We treated clonal *Smo*^-/-^ NIH/3T3 cells expressing the cilia PKA biosensor with FSK and SAG. In *Smo*^-/-^ cells, SAG had no effect on cilia PKA biosensor activity (**Fig. 3E,G**), indicating that SMO is critical for suppressing ciliary PKA activity in response to HH pathway activation.

Cyclopamine (CYA) is a small molecule inhibitor of SMO that triggers accumulation of SMO at the primary cilium but keeps SMO in an inactive conformation (***17, 50, 52–54***). Treatment of cilia PKA biosensor-expressing NIH/3T3 cells with CYA, accordingly, caused SMO to accumulate at primary cilia, at levels similar to that caused by treatment with SAG (**Fig. S3A-E**). Unlike SAG, CYA did not prevent FSK-mediated activation of the cilia PKA biosensor (**Fig. 3D,F**). Thus, SMO localization to the primary cilium is not sufficient to inhibit ciliary PKA; SMO must also be in an active state.

### GRK2/3 phosphorylation of SMO is required to suppress PKA activity at the cilium

G protein-coupled Receptor Kinases 2 and 3 (GRK2/3) promote vertebrate HH signal transduction, and how they regulate HH signal transduction is being actively investigated (***5, 32, 34, 37, 55–57***). For many activated GPCRs, GRKs participate in desensitization (***58***). Indeed, phosphorylation of GPR161 by GRK2 triggers β-Arrestin recruitment and trafficking of GPR161 out of the cilium (***36***). However, GRK2/3 has GPR161-independent functions in HH signaling, as GRK2/3 is required for activation of HH target gene transcription even in the absence of GPR161 (***37***).

GRK2/3 is also able to phosphorylate the C-terminal tail of SMO (***32, 55, 59, 60***), and GRK2/3-phosphorylated SMO is enriched in the primary cilium following HH activation (***60***). One hypothesis is that GRK2/3 mediates the interaction between SMO and PKA by phosphorylating the C-terminal tail of ciliary SMO, allowing SMO to directly bind and inhibit the catalytic subunit of PKA (PKA-C) phosphorylating the C-terminal tail of ciliary SMO, allowing SMO to directly bind and inhibit the catalytic subunit of PKA (PKA-C) (***32, 34, 60***) (**Fig. 4A**). We asked whether SMO-mediated inhibition of ciliary PKA depends on GRK2/3 activity.

**Figure 4:**
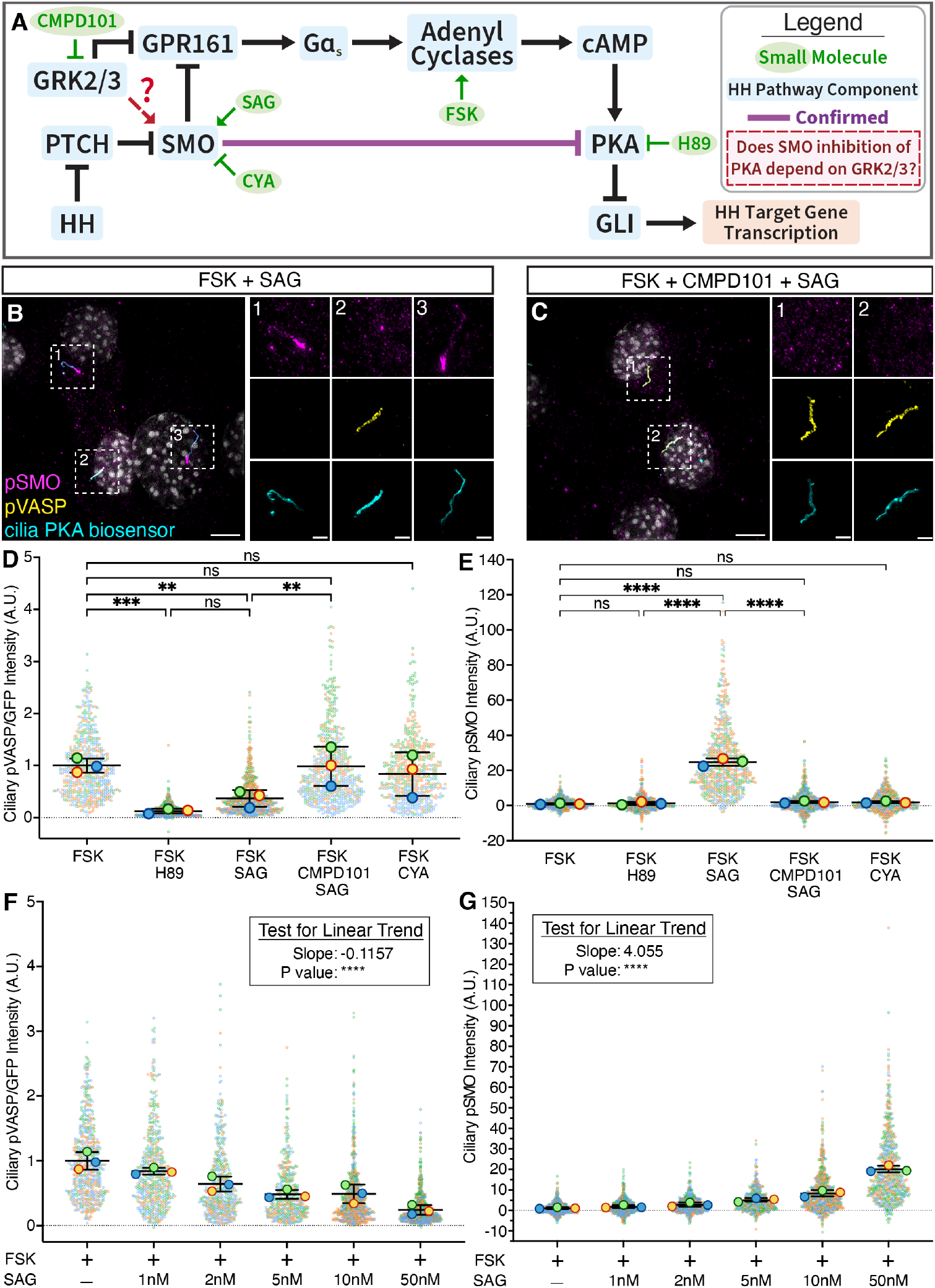
GRK2/3 activity is required for SMO to suppress ciliary PKA activity. (**A**) Schematic of the working model of HH signaling. Red arrow indicates part of the system investigated in this figure. (**B-C**) Immunofluorescence images of cilia PKA biosensor cells. Cells were treated with SAG and FSK (100nM for 24h and100nM for 15 minutes, respectively) (B), or SAG, CMPD101 and FSK (100nM, 30μM and 100nM for 24h, 24h and 15 minutes, respectively) (C). Images depict cells stained for pVASP (pVASPS157, yellow), cilia PKA biosensor (GFP, cyan), pSMO (phospho-SMO S362/S363/S364, magenta) and nuclei (Hoescht, grey). Scale bars for larger images are 5μM, and for insets are 2.5μM. (**D**) Quantification of ciliary pVASP intensity in cilia PKA biosensor cells stained for pVASP. Cells were treated with SAG (100nM), SAG and CMPD101 (100nM and 30μM, respectively), CYA (5μM) or H89 (20μM for 15 minutes) and FSK (100nM for 15 minutes). (**E**) As with D, but for the quantification of ciliary pSMO intensity in the same cells. (**F**) Quantification of ciliary pVASP intensity in cilia PKA biosensor cells. Cells were treated with increasing dosages of SAG (for 24h) and FSK (75nM for 15 minutes). Significance was determined by a one-way ANOVA followed by a post-test for linear trend. (**G**) As with F, but for the quantification of ciliary pSMO intensity in the same cells. For all plots, each biological replicate is color-coded. For D and E, significance was determined via one-way ANOVA of the means of each biological replicate, followed by Šídák’s multiple comparison test (panels D and E), or by a test for linear trend (panels F and G). P values are indicated as follows: ** p < 0.003, ***p < 0.0002, and ****p < 0.0001. Data are represented are means of replicates ± SD.

To test whether GRK2/3 acts at the level of SMO to suppress ciliary PKA during HH signaling, we employed CMPD101, a pharmacological inhibitor of GRK2/3 (***61***). Consistent with previous findings (***60***), activating SMO with SAG increased ciliary SMO phosphorylated at a GRK2/3 consensus site and CMPD101 blocked GRK2/3-dependent phosphorylation of SMO (**Fig. 4B,C**). Treatment of cilia PKA biosensor-expressing NIH/3T3 cells with FSK, SAG and CMPD101 revealed that CMPD101 blocked the ability of SMO to inhibit PKA at the cilium (**Fig. 4B-E**). CYA treatment of cilia PKA biosensor cells neither induced induce SMO inhibition of PKA, nor GRK2/3-dependent phosphorylation of SMO (**Fig. 4F,G**). Thus, GRK2/3 activity is required for SMO to inhibit PKA in cilia.

To assess how SMO activation affects its phosphorylation and ability to inhibit ciliary PKA, we measured the SAG dose response of cilia PKA biosensor activity and ciliary phosphorylated SMO levels. SAG increased SMO phosphorylation and decreased cilia PKA biosensor activity in a dose-dependent way (**Fig. 4F-G**).

Interestingly, immunofluorescence imaging of GRK2/3 phosphorylated SMO (pSMO) and pVASP in cells treated with FSK and intermediate levels of SAG revealed that cells were heterogeneous, exhibiting ciliary pVASP or pSMO, but not high levels of both (**Fig. S4**). Ciliary SMO phosphorylation and ciliary PKA activity were therefore anti-correlated both at the population level and the single cell level. We conclude that GRK2/3 phosphorylation of SMO is critical for inhibition of ciliary PKA and that GRK2/3 and PKA ciliary activity are mutually exclusive.

### Gα_i_ inhibits ciliary PKA activity downstream of a ciliary GPCR, but not downstream of SMO

In addition to GPR161, a growing number of GPCRs have been described that localize to and uniquely function at cilia (**62**). For many of these cilia-localized GPCRs, it is unclear whether they act via PKA or other effectors. To test whether a ciliary GPCR also affects ciliary PKA activity, we created cilia PKA biosensor NIH/3T3 cells stably expressing somatostatin receptor 3 (SSTR3). SSTR3 is a ciliary GPCR which couples to Gα_i/o_ for its downstream signaling (**63–66**) (**Fig. 5A**). Stimulating FSK-treated SSTR3-expressing cells with somatostatin (SST) inhibited cilia PKA biosensor activity (**Fig. 5B, 5D**). Thus, a ciliary GPCR also regulates ciliary PKA activity.

**Figure 5:**
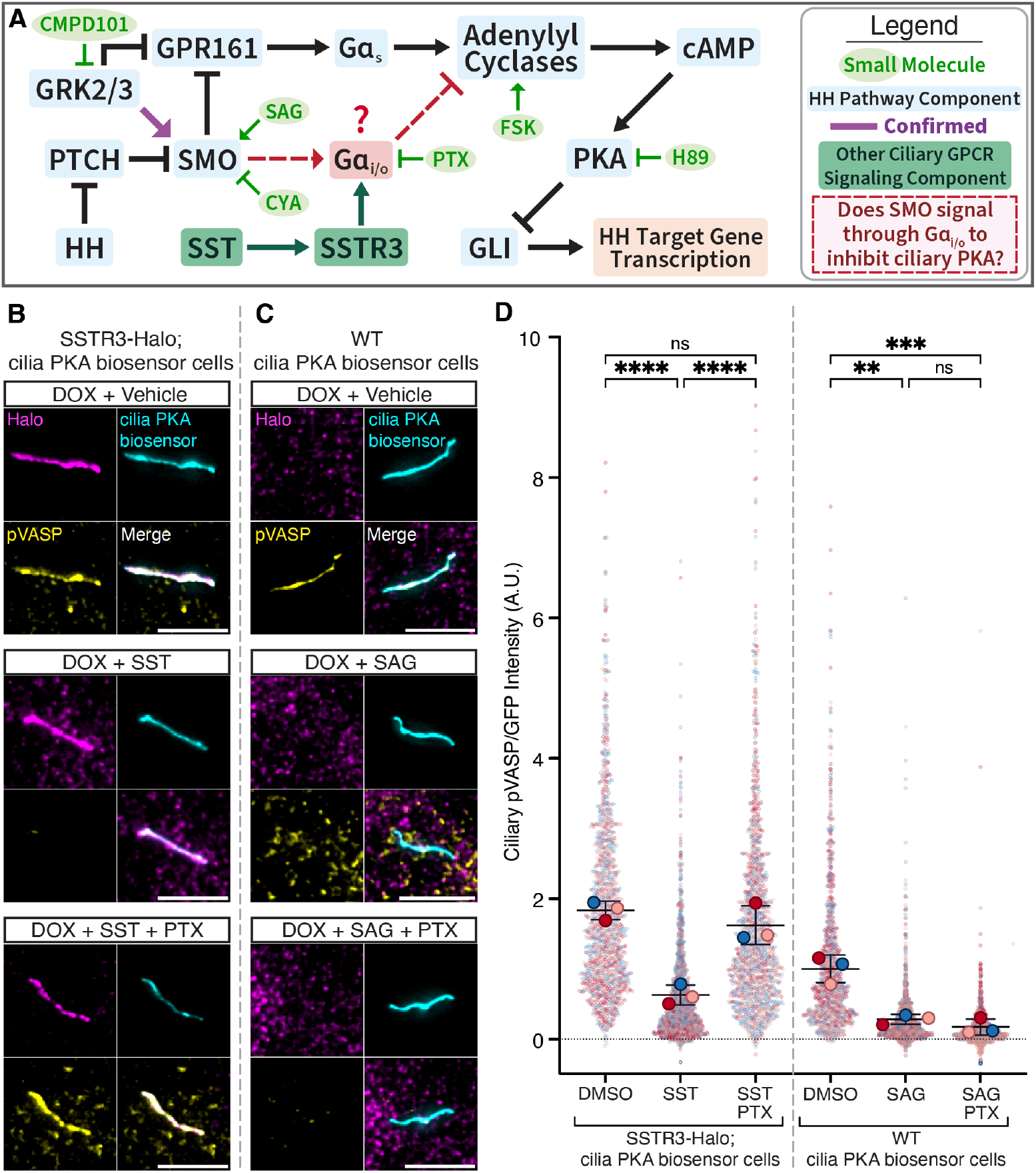
Gα_i_ inhibits ciliary PKA, but does not mediate SMO-based control of ciliary PKA. (**A**) Schematic of the working model of HH signaling. Red arrows indicate areas of inquiry relevant to this figure. (**B**) Immunofluorescence imaging of cilia PKA biosensor cells stably expressing Halo-tagged SSTR3 in response to doxycycline (DOX). Cells were serum starved and treated with either DMSO, SST (10μM for 2h), or SST and PTX (10μM for 2h and 100ng/mL for 16h, respectively). Scale bars are 5μM. (**C**) Immuno-fluorescence imaging of cilia PKA bio-sensor cells. Cells were serum starved and treated with either DMSO, SAG (100nM for 24h), or SAG and PTX (100nM for 24h and 100ng/mL for 16h, respectively). (**D**) Quantification of ciliary pVASP intensity, normalized to ciliary GFP, of C and D. Each biological replicate has its own color. Significance was determined via one-way ANOVA of the means of each biological replicate, followed by Šídák’s multiple comparison test. P values are indicated as follows: **p < 0.003, ***p < 0.0002, and ****p < 0.0001. Data are represented are means of replicates ± SD.

To assay the dependency Gα_i/o_ on the ability of SSTR3 to inhibit ciliary PKA, we treated our cells with pertussis toxin (PTX), an inhibitor of Gα_i/o_ proteins (***67, 68***). The ability of SST to inhibit cilia PKA biosensor activity was blocked by PTX (**Fig. 5B, 5D**), indicating that Gα_i/o_ can control ciliary PKA activity. Interestingly, we previously found that SST activation of SSTR3-expressing fibroblasts induces *Gli1* similarly to SAG, suggesting that activation of ciliary Gα_i/o_ can activate the HH transcriptional response (***14***). We conclude that upon stimulation with SST, SSTR3 can stimulate Gα_i/o_ and inhibit ciliary PKA to drive GLI-mediated HH target gene transcription.

Gα_i/o_ has been investigated as an effector of SMO in HH signal transduction (***26–32***). To test whether Gα_i/o_ is required for SMO to inhibit PKA specifically in the primary cilium, we treated cilia PKA biosensor-expressing NIH/3T3 cells with SAG and PTX. Unlike SST activation of SSTR3, PTX did not block the ability of SAG activation of SMO to inhibit cilia PKA biosensor activity (**Fig. 5B-D**). Thus, despite Gα_i/o_ being critical for ciliary GPCR-mediated control of PKA, Gα_i/o_ activity is dispensable for SMO-mediated inhibition of PKA in the primary cilium.

### SMO A635 contributes to inhibition of ciliary PKA

An alternative to the hypothesis that Gα_i/o_ mediates SMO inhibition of ciliary PKA is that SMO signals by the direct binding and inhibition of PKA-C via a PKI-like motif (***32, 34, 60***). A portion of the SMO proximal carboxy tail (pCT, residues 622–638) resembles PKI motifs of other PKA inhibitory proteins and these residues are necessary for SMO function in zebrafish (***34***).

Typically, PKA-C phosphorylates its substrates at a serine or threonine at a consensus (RRXS/TΦ where Φ is a hydrophobic residue) phosphorylation site (P site) (***69–71***). Pseudosubstrates differ from substrates in having a non-phosphorylatable residue at the P site. Replacement of the P site residue of a PKA pseudosubstrate with serine converts them into substrates and increases dissociation from PKA-C (***71, 72***). Happ et al. previously demonstrated that a version of SMO in which the pCT SMO PKI motif P site alanine was substituted with serine (SMO-A635S) did not activate the HH transcriptional response (***34***).

To assess whether converting the pCT SMO PKI motif to a PKA substrate affects the ability of SMO to inhibit ciliary PKA, we generated *Smo*^-/-^ cilia PKA biosensor NIH/3T3 cells stably expressing HALO-tagged SMO-A635S and stimulated them with SAG (**Fig. 6A**). Compared to wild-type SMO, SMO-A635S exhibited attenuated inhibition of cilia PKA biosensor activity (**Fig. 6B-C**). Unlike wild-type SMO, SMO-A635S did not induce *Gli1* in response to SAG (**Fig. 6D**). Thus, the PKI motif in SMO is critical for its ability to control ciliary PKA activity and activate the downstream pathway.

**Figure 6:**
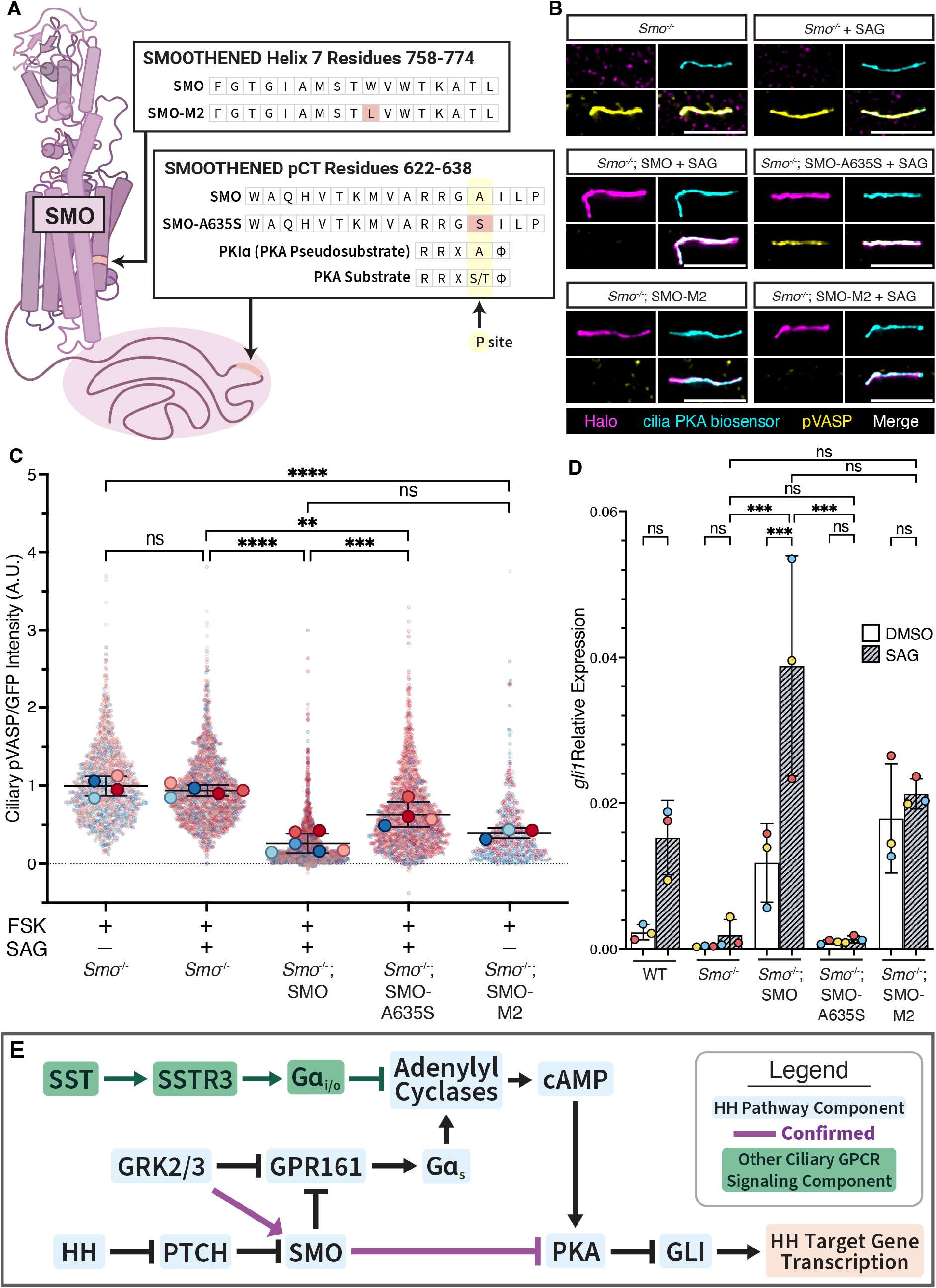
SMO PKI motif contributes to inhibiting ciliary PKA. (**A**) A schematic of the SMO mutations assessed in this figure. (**B**) *Smo*^-/-^ cilia PKA biosensor cells, stably expressing Halo-tagged wild-type SMO, SMO-A635S or SMO-M2, as indicated. Cells were treated SAG (100nM for 24h) and FSK (100nM for 15 minutes). Scale bars are 5μM. (**C**) Quantification of ciliary pVASP intensity, normalized to ciliary GFP. Significance was determined via one-way ANOVA of the means of each biological replicate, followed by Šídák’s multiple comparison test. (**D**) qRT-PCR of *Gli1* in wild-type or *Smo*^-/-^ cells expressing wild-type SMO, SMO-A635S or SMO-M2 and treated with SAG, as indicated. Significance was determined via two-way ANOVA of the means of each biological replicate, followed by Šídák’s multiple comparison test. (**E**) Schematic of HH signaling. For all plots, each biological replicate is color coded. P values are indicated as follows: ** p < 0.003, ***p < 0.0002 and ****p < 0.0001. Data are represented are means of replicates ± SD.

A constitutively active, oncogenic single amino acid substitution in SMO W5535L, better known as SMO-M2, is sufficient to cause basal cell carcinoma, medulloblastoma and rhabdomyosarcoma (***73***). To assess whether oncogenic mutations affect the ability of SMO to inhibit ciliary PKA, we generated *Smo*^-/-^ cilia PKA biosensor NIH/3T3 cells stably expressing HALO-tagged SMO-M2. Unlike wild-type SMO, SMO-M2 inhibited cilia PKA biosensor activity even in the absence of SAG (**Fig. 6B-C**). Similarly, SMO-M2 induced *Gli1* even in the absence of SAG (**Fig. 6D**). Thus, an oncogenic mutation constitutively activates the ability of SMO to inhibit ciliary PKA.

To assess whether any of these mutations uncovered a cryptic dependency on Gα_i/o_ we treated each of the mutant SMO-expressing cells with SAG and PTX. PTX did not attenuate the ability of any mutant SMO to inhibit cilia PKA biosensor activity, consistent with Gα_i/o_ being dispensable for SMO signaling (**Fig. S6**). We conclude that SMO signals independently of Gα_i/o_. Rather, the SMO pseudo-substrate site-mediated inhibition of PKA activity in the cilium activates the downstream HH signal transduction pathway (**Fig. 6E**).

## Discussion

Although vertebrate SMO requires the cilium to activate the downstream HH signal transduction pathway, how it does so has been elusive. Recent revelatory work has discovered that the C-terminus of vertebrate SMO can bind and inhibit PKA (***32, 34***). PKA phosphorylates GLI to inhibit HH target gene transcription (***18–20***). Because SMO, PKA and GLI all can localize to primary cilia (***14, 16, 18, 60***), we investigated whether SMO signals through inhibiting PKA at the primary cilium. To test this hypothesis, we developed a biosensor that measures ciliary PKA activity.

This biosensor revealed that SMO inhibits ciliary PKA in a state-dependent manner. SHH, (which acts through its receptor PTCH1 to activate SMO) (***74***) and SAG (which directly binds and activates SMO) (***50, 51***) both induced SMO to accumulate at cilia and inhibit ciliary PKA. In contrast, CYA, a SMO inhibitor (***52, 75***), induced SMO to accumulate at cilia but not inhibit ciliary PKA. Thus, the accumulation of SMO in cilia is not sufficient to inhibit ciliary PKA; SMO must also be in an active state.

Another requirement for SMO to inhibit ciliary PKA is GRK2/3: SMO is phosphorylated by GRK2/3 (***60***) and pharmacological inhibition of GRK2/3 prevented SMO from inhibiting ciliary PKA. Consistent with this conclusion, CYA prevented GRK2/3 from phosphorylating SMO and from inhibiting ciliary PKA. GRK2/3 also phosphorylates GPR161, facilitating the removal of GPR161 from the primary cilium (***36***), but GRK2/3 can still regulate HH signaling in the absence of GPR161 (***37***). Thus, another way in which GRK2/3 participates in HH signal transduction is by phosphorylating SMO to suppress ciliary PKA.

By measuring ciliary PKA activity, we assessed molecular mechanisms by which SMO could control PKA. Two models have been: 1) SMO activates Gα_i/o_ to inhibit production of cAMP, thereby inhibiting PKA (***26–30***), and 2) SMO acts like a PKI to directly bind and inactivate PKA (***32, 34***). We found that Gα_i_ activity is dispensable for SMO to inhibit PKA. Furthermore, a mutation predicted to turn the PKI-like motif into a PKA substrate (and thus prevent PKA inhibition) blocked the ability to SMO to inhibit PKA and activate downstream transcription. As PKA substrates are generally dissociated from PKA upon phosphorylation (***71, 72***), we propose that converting SMO from a pseudosubstrate to a substrate increases its dissociation rate from PKA-C and disrupts its ability to inhibit PKA.

Additionally, we find that an oncogenic form of SMO constitutively inhibits ciliary PKA activity. Oncogenic mutations in SMO cause a number of cancers, including medulloblastoma, basal cell carcinoma and rhabdomyosarcoma (***76***). Our findings suggest that, for SMO, constitutive ciliary localization, constitutive ciliary PKA inhibition and oncogenic activity are causally related. Although HH pathway-related medulloblastoma is responsive to small molecule inhibitors of SMO, acquisition of SMO mutations that interfere with drug binding lead to recurrence (***77, 78***). We propose that an alternative therapeutic approach that bypasses SMO is to reactivate ciliary PKA.

Interestingly, SSTR3, a ciliary Gα_i_-coupled GPCR not implicated in regulating HH signaling, could also inhibit ciliary PKA and activate HH transcription when expressed in fibroblasts. As the primary cilium is a specialized microenvironment for multiple receptors and signaling pathways, how many are mediated via ciliary PKA? A burgeoning number of other GPCRs have more recently been found to localize to primary cilia across many different tissues (***35, 79–83***). For example, melanocortin 4 receptor (MC4R) is a neuronal cilium-localized GPCR critical for the control of feeding behavior and long-term energy homeostasis (***83–86***). Similarly, SSTR3 endogenously localizes to the primary cilia of mammalian neurons and is implicated in learning and memory (***63, 64***). One possibility is that there are different effectors other than PKA for other ciliary GPCRs. Another possibility is that many ciliary GPCRs act through ciliary PKA but achieve cell type-specific effects via different effectors downstream of PKA. In the case of SSTR3, it is unclear if SSTR3-expressing cells also endogenously express GLI proteins. Thus, it is possible that ciliary PKA phosphorylates a different effector in SSTR3-expressing cells.

Moreover, it is unclear if cells simultaneously communicate via multiple ciliary signaling pathways, such as the HH signal transduction pathway and ciliary GPCR signaling pathways. It is possible that the set of cells competent for HH signaling and the set of cells that are competent for other ciliary GPCR signaling are mutually exclusive, thus preventing crosstalk within their cilia. Alternatively, all Gα_i/o_- or Gα_s_-coupled ciliary GPCRs in cells competent for HH signaling may influence GLI-dependent transcription. Untangling GPCR signaling at the primary cilium will be critical to understanding, not only mechanisms of HH signaling, but how a host of signals critical to human health are transduced.

## Acknowledgements

We thank Roshanak Irannejad for the VASP plasmid; Aaron Marley for the SSTR3 plasmid; members of the Reiter laboratory for discussion and advice; Licia Selleri, Tien Peng, and Frances Ding for comments on the manuscript; DeLaine Larsen, Kari Herrington, SoYeon Kim, Micaela Lasser, and Nico Stuurman from the UCSF Center for Advanced Light Microscopy for microscope use and imaging assistance; Sara Elmes from the UCSF Laboratory for Cell Analysis for cell sorting instrument use and flow cytometry; Gary Moulder, Louie Ramos, William Figueroa, Francesca Penny, and Stephanie Gilbert from the UCSF Cardiovascular Building fish facility for fish husbandry; and Frances Ding for assistance writing data processing scripts.

This paper was typeset with the bioRxiv word template by @Chrelli: www.github.com/chrelli/bioRxiv-word-template

## Funding

This work was also supported by a grant from the NIH (R01AR054396) to J.F.R. Data for this study were acquired at the Center for Advanced Light Microscopy at UCSF on an OMX-SR obtained using grants from the NIH (5R35GM118119), the UCSF Program for Breakthrough Biomedical Research funded in part by the Sandler Foundation, the UCSF Research Resource Fund Award, and HHMI. Flow cytometry was performed at the UCSF Helen Diller Family Comprehensive Cancer Center LCA and LCA-Genomic Core Facility using grants from the NIH (P30CA082103).

## Author contributions

Conceptualization: TDN, JFR

Methodology: TDN, JFR

Investigation: TDN, MJK, LDC

Visualization: TDN

Funding acquisition: JFR

Supervision: JFR

Writing – original draft: TDN

Writing – review & editing: TDN, JFR

## Competing interest statement

J.F.R. cofounded Renasant Bio and a company funded via BridgeBio.

## Materials and Methods

### Vector construction and generation of stable cell lines

To generate NIH/3T3 Flp-In cell lines expressing the ciliary PKA biosensor, ARL13B-GFP-VASP was cloned with the In-Fusion HD cloning kit (Takara, 639650) into a version of pGLAP5 with an attenuated EF1a promoter lacking the TATA box (***87***), a backbone previously generated in our lab (***14***). We transfected cells with this plasmid, concurrently with the pOG44 Flp-Recombinase Expression Vector (Invitrogen, V600520), with Lipofectamine LTX (Invitrogen, 15338100), according to the ThermoFisher Flp-In System protocol to generate stable Flp-In expression cell lines. Cells were selected with 70μg/mL of hygromycin B (Corning, 30-240-CR). Following selection, we selected a single clone of these cells to characterize and build all subsequent cell lines. Plasmid encoding VASP amino acids 148-164 was a generous gift from Roshanak Irannejad.

For generating mRNA encoding ARL13B-GFP-VASP, ARL13B-GFP-VASP was cloned into the pCS107 expression vector using the In-Fusion HD cloning kit (Takara, 639650).

To generate cell lines expressing Halo-tagged SSTR3, SMO, SMO-A635S, and SMO-M2, we cloned each protein of interest into a pLVX-TetOne-Puro backbone (Takara 631849).

We used clustered regularly interspaced short palindromic repeats (CRISPR)-mediated editing to generate loss-of-function mutations in Smo using two different guide RNAs per gene in our cilia PKA biosensor cells. Synthetic guide RNAs for *Smo* (5’-CCCACGCACGGGGCGGCCAG-3’, 5’-UCCCGCUCAAGGCCGCCCCC-3’) were ordered from Synthego, complexed with TrueCut Cas9 Protein v2 (Invitrogen, A36496), and nucleofected with the Neon Transfection System (Invitrogen). Cells were clonally selected and screened via PCR for genomic deletions with primers for the genomic regions of interest for mouse *Smo* (forward primer 5’-AG-GGTTCCCAGGGTTGAAGA-3’, reverse primer: 5’ -CACACGTTGTAGCG-CAAAGG-3’).

Lentivirus was generated by transfecting 7.5 μg of each plasmid of interest with 1.5 μg of pCMV-VSV-G (Addgene, 8454) and 6 μg of psPAX2 (Addgene, 12260) into a 10 cm plate of Lenti-X 293T cells (Takara, 632180) at 70–80% confluence using Fugene 6 transfection reagent (Promega, E2691). Medium containing lentiviral particles was collected 1 day after transfection and was concentrated with a Lenti-X Concentrator (Takara, 631232), incubated overnight at 4°C, and followed by centrifugation at 4000g for 45 min at 4°C. The pellet was resuspended in 120μL DPBS (Gibco, 14-90-250).

To generate SMO expression cell lines, cilia PKA biosensor *Smo*^-/-^ cells were transduced with 20μL of resuspended lentivirus containing either doxycycline-inducible SMO-Halo, SMO-A635S-Halo, SMO-M2-Halo in the presence of 4 μg/mL polybrene. 24 hours after transduction, cells were selected with 1μg/mL puromycin (Gibco, A11138-03) for 5 days. Cells that survived selection where then incubated with 100ng/mL doxycycline (Sigma-Aldrich, D5207) for 48 hours and incubated in HaloTag Alexa Fluor 660 Ligand (Promega, G8472) overnight before being enriched via fluorescence-activated cell sorting (FACS) on a BD FACSAria III Cell Sorter.

To generate SSTR3 overexpression cell lines, cilia PKA biosensor cells were transduced with lentivirus containing doxycycline-inducible SSTR3-Halo-puro in the presence of 4 μg/mL polybrene. Cells were then transduced with 1μg/mL doxycycline (Sigma-Aldrich, D5207) for 48 hours, selected with 1μg/mL puromycin (Gibco, A11138-03). Cells were then incubated in HaloTag Alexa Fluor 660 Ligand (Promega, G8472) overnight before being enriched via fluorescence-activated cell sorting (FACS) on an BD FACSAria III Cell Sorter.

## mRNA synthesis

To generate mRNA for expressing the cilia PKA biosensor in zebrafish, we grew pCS107-ARL13B-GFP-VASP in *dam*^*–*^*/dcm*^*–*^ competent *E. coli* (New England Biolabs, C2925H). We isolated our plasmid with the Plasmid Plus Midi Kit (QIAGEN, 12943). For zebrafish injections, we linearized the construct with *ApaI* (New England Biolabs, R0114L) and generated mRNA with the mMESSAGE mMACHINE SP6 kit (Invitrogen, AM1340).

## Zebrafish husbandry and mRNA injection

Adult *Danio rerio* zebrafish were maintained under standard laboratory conditions. Zebrafish of Ekkwill (EKW) background were used as wild type. Embryos were maintained in egg water containing 60 μg/mL sea salt (Instant Ocean) in distilled water. We injected 500pg of ARL13B-GFP-VASP mRNA at the one-cell stage. We incubated injected embryos in egg water, and unfertilized embryos were removed 6 hours post injection. All zebrafish protocols were approved by the Institutional Animal Care and Use Committee (IACUC) of the University of California, San Francisco.

### Mammalian cell culture

NIH/3T3 Flp-In cells (Invitrogen, R761-07) were cultured in Dulbecco’s modified Eagle’s medium with high glucose (Gibco, 11965118) supplemented with 10% newborn calf serum (Gibco, 16010159) and GlutaMAX supplement (Gibco, 35050061). Cells were treated with antibiotic-antimycotic (Gibco, 152400632) following FACS for 1 week. Cells were otherwise maintained in the absence of antibiotics. To induce ciliation, cells were grown to confluence and starved overnight in Opti-MEM reduced serum medium with GlutaMAX supplement (Gibco, 51985091).

### Immunofluorescence staining

We seeded cells on 12 mm cover glasses of 170μm thickness (Paul Marienfeld, 0117520) at a density of 8×10^4^ cells per well in a 24-well plate. After drug treatment and induction of ciliation, we fixed cells for 10 minutes in 4% PFA (VWR, 100504-782) diluted in DPBS (Gibco, 14-90-250). We diluted primary antibodies in blocking buffer (0.1% TritonX-100, 0.02% Sodium Azide, 3% BSA) and incubated them overnight at 4°C. Subsequently we incubated cells in donkey Alexa Fluor-conjugated secondary antibodies, ChromoTek GFP-Booster Alexa Fluor 488 (Proteintech, gb2AF488), and Hoechst (ThermoFisher Scientific) diluted in blocking buffer at room temperature for 1 hour. We mounted cover glasses in ProLong Glass antifade mountant (Invitrogen, P36982) and allowed the slides to cure overnight at room temperature before imaging.

We fixed dechorionated zebrafish embryos in 4% PFA (VWR, 100504-782) diluted in DPBS (Gibco, 14-90-250) for 2 hours at room temperature. We blocked embryos in 1% BSA, 1% DMSO and 0.5% Triton X-100 in PBS (PBDT) for 1 hour. After blocking, we incubated embryos overnight at 4°C with primary antibodies diluted in PBDT. Subsequently we incubated embryos in donkey Alexa Fluor-conjugated secondary antibodies, ChromoTek GFP-Booster Alexa Fluor 488 (Proteintech, gb2AF488), and Hoechst (ThermoFisher Scientific) diluted in PBDT for 2 hours at room temperature. We incubated embryos in 70% glycerol overnight, then mounted in in ProLong Glass antifade mountant (Invitrogen, P36982) and allowed the slides to cure overnight at room temperature before imaging.

### Image Acquisition and Ciliary Fluorescence Intensity Quantification

We imaged fixed cells on a DeltaVision-OMX-SR (GE Healthcare) equipped with a Plan ApoN 60X/1.42 Oil objective and three PCO.edge 5.5 15bit sCMOS Cameras (liquid cooled). Four-channel fluorescence imaging was captured with a Toptica 4 line laser launch light source, laser excitation wavelengths 405nm/488nm/568nm/642nm, and emission filters 435/31m, 528/48m, 609/37m, and 683/40m. Images for quantification were acquired using the widefield setting, and representative images, where indicated in the figure legend, were acquired with 3D-SIM Data. Immersion oil with refractive index of 1.518 was used for most experiments. Z stacks of 5–6 μm were collected using a 0.250μm step size for widefield imaging and 0.125 μm step size for 3D-SIM imaging. Raw images were reconstructed using SoftWorx 6.5.2 (GE Healthcare) using default parameters.

We imaged fixed zebrafish embryos with Zeiss LSM 800 laser scanning confocal microscope equipped with a 63x/1.4 oil immersion objective and captured using the Zen Imaging Software (Zeiss). While collecting images, we held constant the gain, offset and laser power for each antibody combination. We processed images identically and used ImageJ/FIJI software (***88***) to generate sum and maximal projections.

We used Cell Profiler image analysis software (***89***) on our sum projection images to generate fluorescence intensity quantifications. A cilia marker (ARL13B-GFP-VASP) was used to identify cilia and create a mask using the object identification module in CellProfiler using differences in signal intensity and size to segment cilia. The ciliary mask was then dilated by 10 pixels to create a dilated ciliary mask. We determined the fluorescence intensity (integrated intensity) and area (in pixels) for both the cilia mask and dilated cilia mask. We determined background-subtracted, area-normalized ciliary fluorescence intensity for all channels of interest with the following formula:

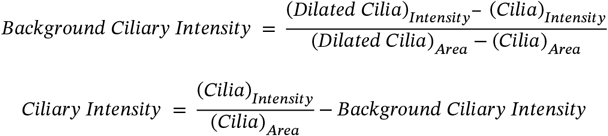

To represent ciliary PKA activity, we report 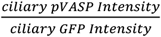 to account for the amount of VASP peptide able to be phosphorylated at the primary cilium in our cilia PKA biosensor cells. Ciliary intensity calculations were ultimately done through a Jupyter Notebook python script. Data were exported to .csv files and graphs were generated in GraphPad Prism 10. Statistical analyses were performed using GraphPad Prism 10. Statistical tests used for each experiment are listed in the accompanying figure legend.

### Quantitative RT-PCR

3T3 cells were seeded in 6-well plates at a density of 300,000 cells per well. Total RNA was extracted using the RNeasy Mini Kit (QIAGEN Cat# 74106) according to the manufacturer’s instructions. cDNA was reverse transcribed from 1ug of RNA using the iSCRIPT cDNA synthesis kit (Bio-Rad Cat#1708891BUN). Each qRT-PCR reaction was performed in technical quadruplicates on a 384 well plate (USA Scientific, Cat# 1438-4700) using PowerUp SYBR Green Master Mix (Applied Biosystems Cat# A25777) and run on a QuantStudio 5 real-time PCR system (Applied Biosciences) running QuantStudio Design & Analysis software (v.1.5.1). We used the following primer sequences: *Hprt* (Forward primer: 5’-CATAAC-CTGGTTCATCATCGC-3’, Reverse primer: 5’-TCCTCCTCAGACCGCTTT T-3’) and *Gli1* (Forward primer: 5’-GGTGCTGCCTATAGCCAGTGTCCTC-3’, Reverse primer: 5’-GTGCCAATCCGGTGG AGTCAGACCC-3’) Relative expression was calculated using the ΔΔCT method normalized to the expression of the housekeeping gene *hprt*.

**Figure S1:**
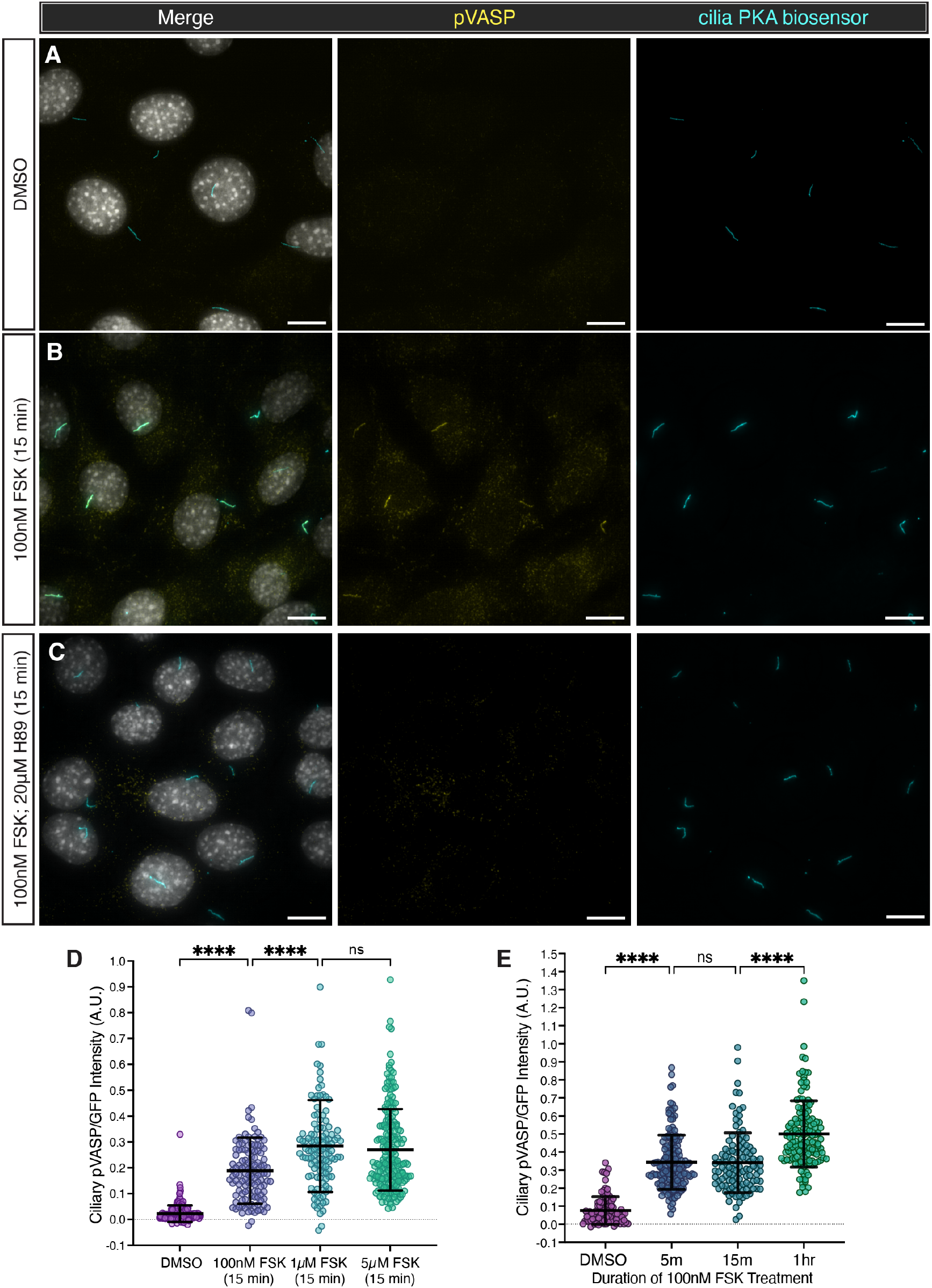
Characterizing the dynamic range of the cilia PKA biosensor. (**A-C**) Representative images of immunofluorescence staining of NIH/3T3 cells stably expressing the cilia PKA biosensor. Cells were serum-starved and then treated with either DMSO, FSK (100nM for 15 minutes), or both FSK and H89 (100nM and 20μM, respectively, for 15 minutes). Images depict cells stained for pVASP (pVASP^S157^, yellow), cilia PKA biosensor (GFP, cyan), and nuclei (Hoescht, grey). Scale bar, 10μM. (**D**) Quantification of ciliary pVASP intensity normalized to ciliary GFP intensity of cells treated with different concentrations of FSK. (**E**) Quantification of ciliary pVASP intensity normalized to ciliary GFP intensity cells treated with FSK for different durations. Significance was determined via one-way ANOVA followed by Tukey’s multiple comparison test. (****p < 0.0001)

**Figure S3:**
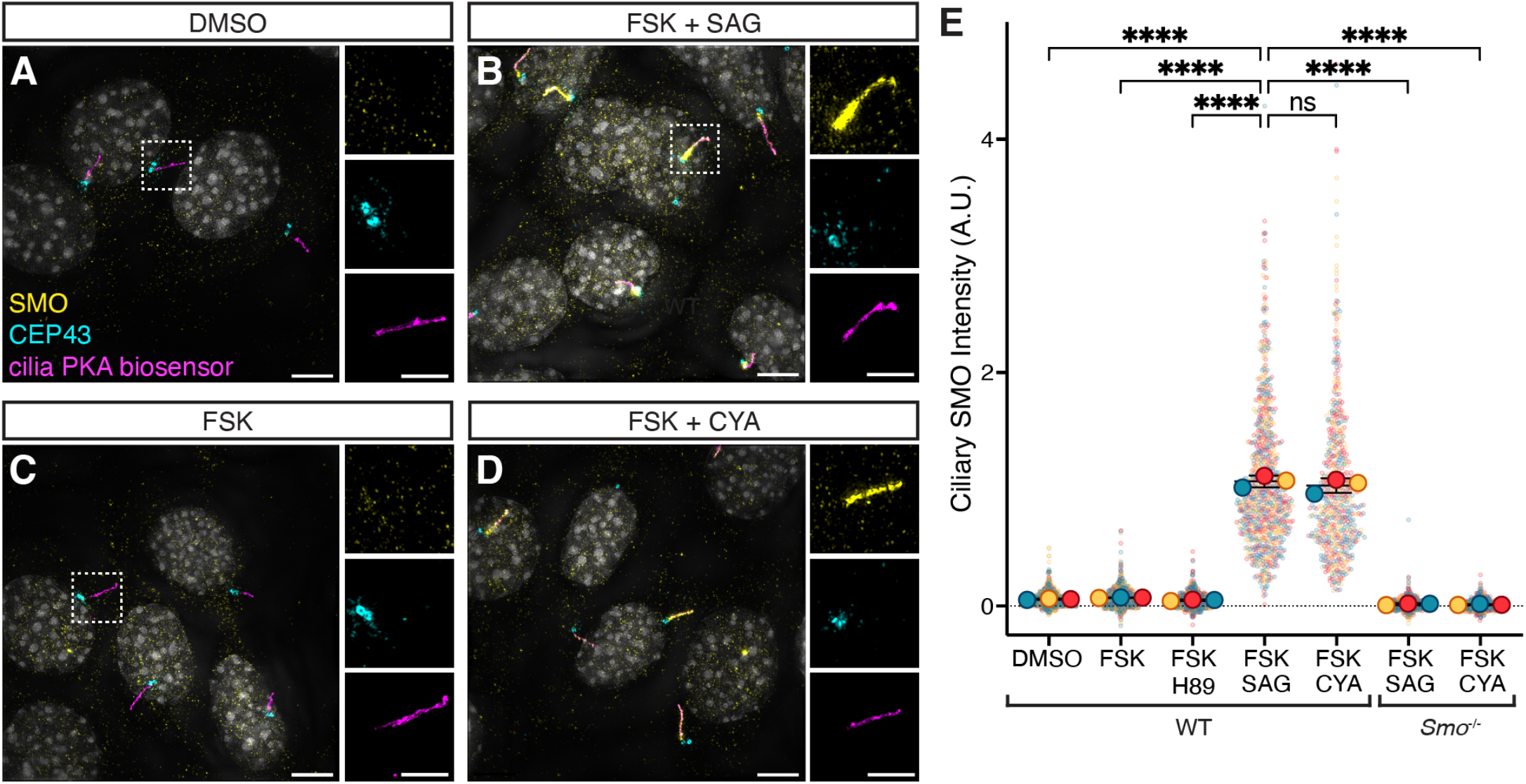
SAG and cyclopamine induce equivalent levels of SMO accumulation at primary cilia. **(A-D**) Immunofluorescence images of cilia PKA biosensor cells treated with the same regimes as in Fig. 3B-G. Images depict cells stained for SMO (pVASP^S157^, yellow), basal bodies (CEP43, cyan), cilia PKA biosensor (GFP, magenta), and nuclei (Hoescht, grey). Scale bars for larger images, 5 μM. Scale bars for insets are 2.5 μM. (**E**) Quantification of ciliary SMO localization from A-D.For all plots, each biological replicate is color coded. Significance was determined via one-way ANOVA of the means of each biological replicate, followed by Šídák’s multiple comparison test. ****p < 0.0001. Data are represented are means of replicates ± SD.

**Figure S4:**
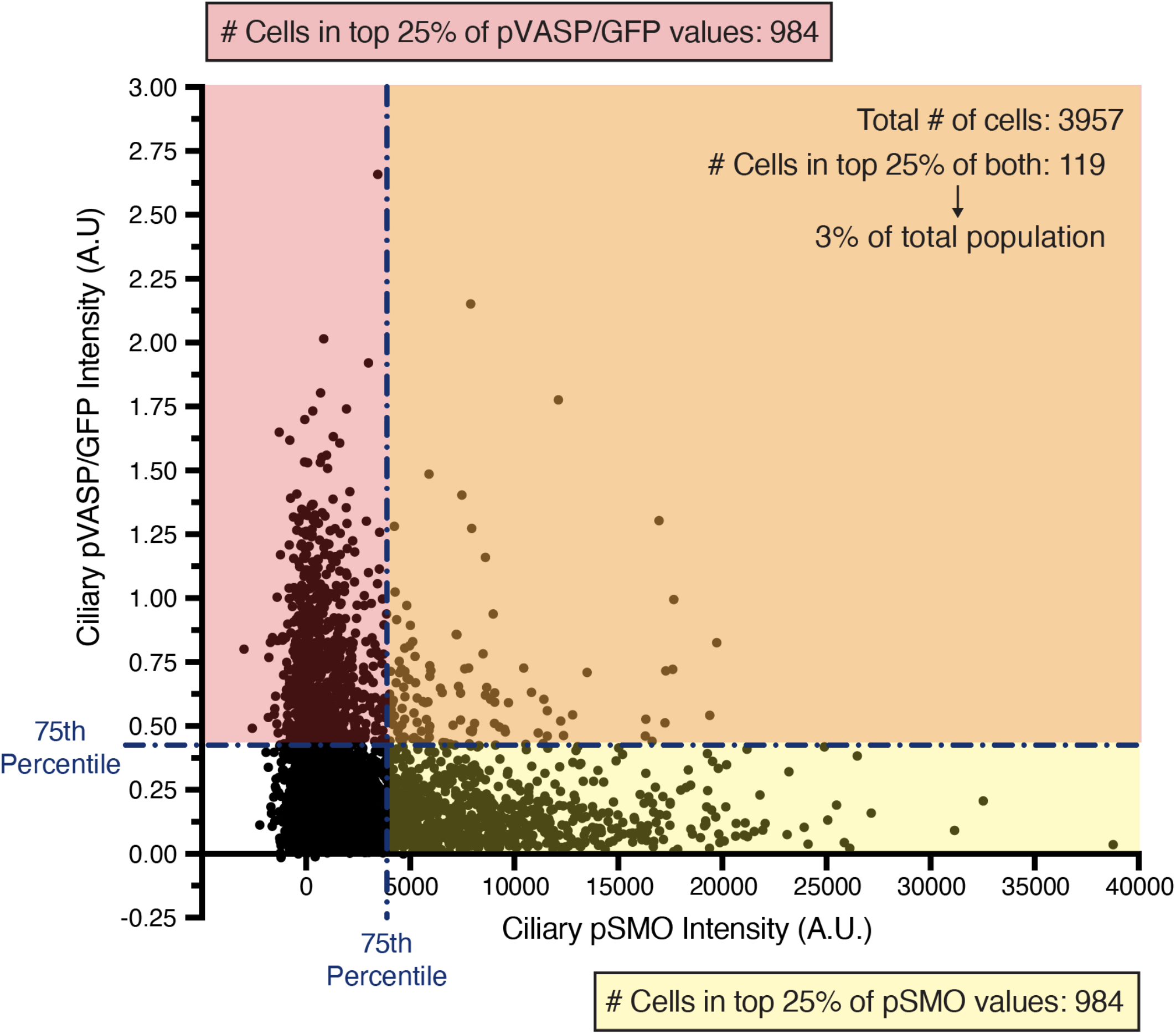
High ciliary pVASP and high ciliary pSMO are mostly mutually exclusive. Each dot represents a cilium of a cilia PKA biosensor cell treated with SAG for 24hrs followed by 15min of 75nM FSK. These data are also used in Figure 3F-G. The x-axis represents the level of ciliary pSMO in each cilium, and the y-axis represents the level of ciliary pVASP/GFP in that same cilium.

**Figure S6:**
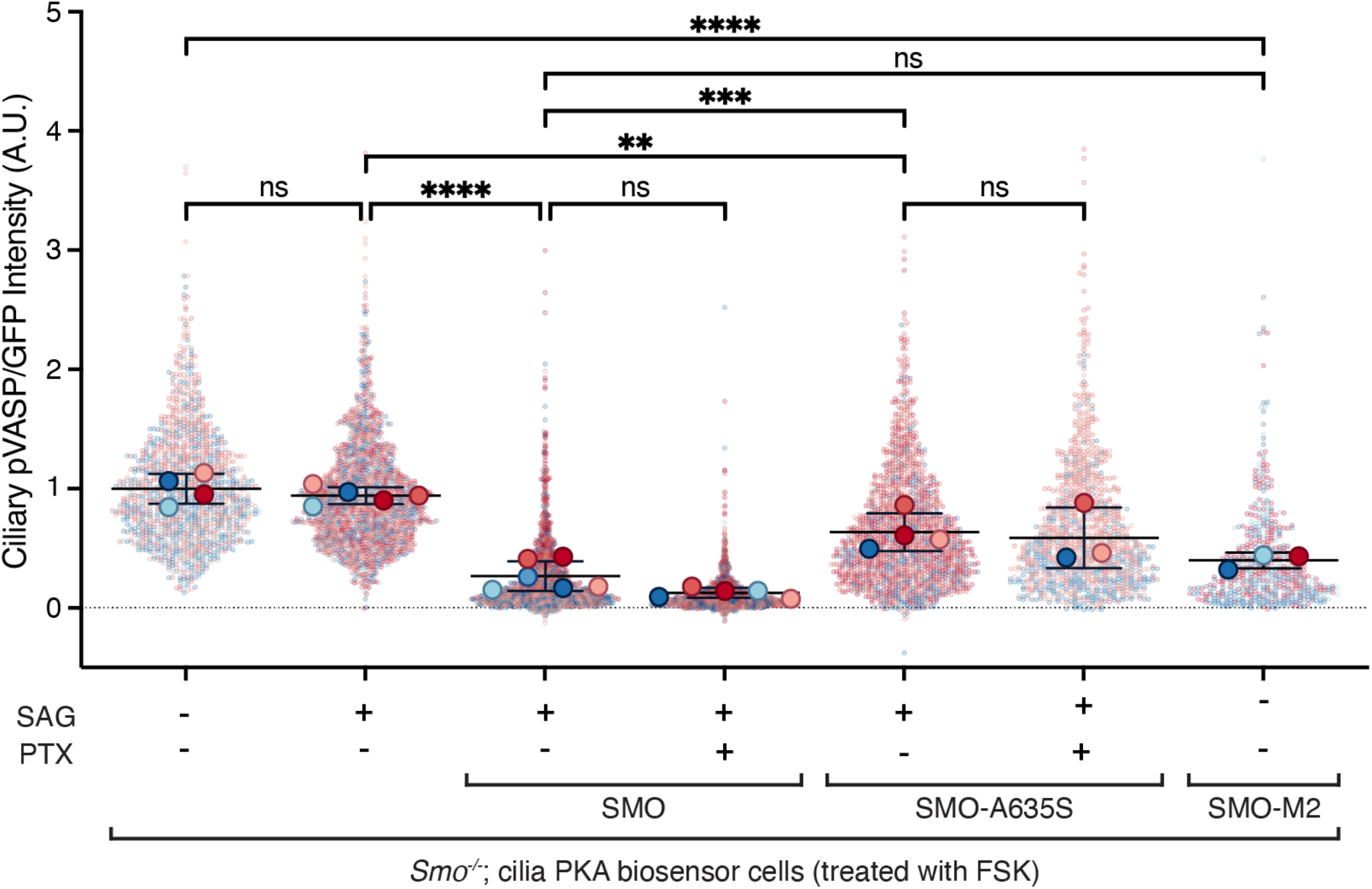
Gα_i/o_ does not synergize with the SMO PKI motif to inhibit ciliary PKA. Quantification of ciliary pVASP intensity, normalized to ciliary GFP, in *Smo*^-/-^ cilia PKA biosensor cells expressing wild-type SMO, SMO-A635S or SMO-M2, as indicated. As indicated, cells were treated SAG (100nM for 24h), or SAG and PTX (100nM for 24h and 100ng/mL for 16h, respectively). All conditions were treated with FSK (100nM for 15 minutes). For all plots, each biological replicate is color coded. Same data as used in Figure 6. Significance was determined via one-way ANOVA of the means of each biological replicate, followed by Šídák’s multiple comparison test. P values are indicated as follows: ** p < 0.003, ***p < 0.0002 and ****p < 0.0001. Data are represented are means of replicates ± SD.

## Notes

### Summary of Updates

Author name corrected to Lorenzo M. Del Castillo.

